# Shared genetic foundation of the DHEAS pathway indicate similar extended childhoods in Neanderthals and modern humans

**DOI:** 10.64898/2026.06.25.734583

**Authors:** Nicholas R. Hartman, Fernando A. Villanea

## Abstract

Adrenarche, the pre-pubertal rise in adrenal androgens, particularly dehydroepiandrosterone (DHEA) and its sulfated form DHEAS, is a critical driver of middle childhood cognitive and social development in modern humans. Compared to other apes, modern human adrenarche is more prolonged, with higher DHEAS levels. Whether this uniquely prolonged human adrenarche is a derived trait of *Homo sapiens* or has deeper hominin roots remains unresolved. Here, we examine the Neanderthal genetic variation in five key DHEAS biosynthesis genes (*HSD3B2, CYP17A1, POR, CYB5A, SULT2A1*). We also examine archaic introgression in these genes by comparing high-coverage Neanderthal genomes with globally diverse modern human sequences from the 1000 Genomes Project. We identify 29 Neanderthal-derived single nucleotide variants across these genes. Key steroidogenic genes in the biosynthesis pathway show no evidence of introgression, consistent with selection on pleiotropic regulators of steroidogenesis. In contrast, accessory genes carried introgressed Neanderthal haplotypes at moderate frequencies in non-African human populations, indicating Neanderthal variants are compatible with the human DHEAS synthesis pathway. All Neanderthal-specific variants were in non-coding regions, with three variants associated with reduced enzyme efficiency or DHEAS production in adults. Additionally, for all 29 positions, the modern human major allele is ancestral, and there is no evidence for a suite of novel adrenarche-extending variants. We conclude that the genetic foundation for extended adrenarche is shared between *Homo sapiens* and Neanderthals, and may have deeper hominin roots. Any phenotypic variation in adrenarche between modern humans and Neanderthals is more likely attributable to differential gene expression than to divergence in protein-coding sequences.

## Introduction

Life history theory is an evolutionary framework that considers how organisms allocate energy among growth, reproduction, and survival, to create diverse developmental strategies (Stearns, 1976). In great apes, extended pre-adult periods evolved under selective pressures favoring slow development, learning, and high parental investment (Jones, 2011). In modern humans this pre-adult period is even more prolonged and includes four stages: infancy, early childhood, middle childhood, and adolescence, each marked by characteristic changes in growth, dentition, cognition, social behavior, and hormones (Bogin, 1997).

For modern humans, middle childhood is a period of major cognitive, social, and emotional transformation (Bogin, 1997; Campbell, 2006). This extended period allows children to acquire cultural knowledge and social skills under parental care, and has been argued to be a key evolutionary innovation contributing to human ecological and cultural success (Bogin, 1997). Yet it remains unclear whether the human-like pattern of middle childhood, closely tied to adrenarche, originated with *Homo sapiens* or has deeper hominin roots. Neanderthals (*Homo neanderthalensis*), our closest extinct relatives, provide a valuable comparison group. The availability of multiple Neanderthal genomes makes it possible to compare the genetic mechanisms that may contribute to variation in adrenarche between Neanderthals and modern humans.

Much of middle childhood cognitive development is driven by adrenarche, a pre-puberty developmental process beginning around age six. It is marked by the expansion of the zona reticularis (ZR) of the adrenal cortex and a corresponding rise in adrenal androgens, specifically dehydroepiandrosterone (DHEA) and its circulating sulfate form, DHEAS (hereafter collectively referred to as DHEA/S) (Nguyen & Conley, 2008). DHEA/S has been hypothesized to affect brain regions involved in social function, memory, complex problem solving, and synaptic development, suggesting a key role in the emergence of human social and cognitive complexity (Campbell, 2006). Increasing DHEA/S also contributes to physical changes such as the growth of axillary and pubic hair and increased muscle and bone mass. (Parker, 1991). Overall, prolonged adrenarche is critical for human social and cognitive development, preparing children for puberty and adulthood.

Because adrenarche, as a hormonal process, leaves no direct trace in fossil material, developmental trajectories in extinct hominins must be reconstructed indirectly from the fossil record. Developmental timing can be inferred from dental growth, which in living primates closely tracks life-history transitions such as adrenarche. In modern humans, the first molar (M1) erupts around age six, coinciding with the onset of middle childhood and adrenarche (Dean, 2006). In chimpanzees (*Pan troglodytes*), M1 emerges earlier, at roughly 3.5 years, corresponding to the start of their adrenarche (B. H. Smith, 1994). Multiple studies indicate that M1 emergence in *Australopithecus* and early *Homo* more closely resembled chimpanzees than modern humans (Dean et al., 2001; Dean & B. H. Smith, 2009; Kelley & Schwartz, 2012; R. Smith et al., 1995; Thompson & Nelson, 2011). These findings support the view that the distinctly prolonged modern human pattern of development had not yet emerged in early *Homo*.

For later *Homo*, fossil-based estimates of life history stages are conflicting. Bermúdez de Castro et al. (1999) argued that *Homo heidelbergensis* showed modern human-like dental growth, but a subsequent study reported faster dental development in the same species (Ramirez Rozzi & Bermúdez de Castro, 2004). Work on Neanderthal ontogeny is likewise mixed, though most studies suggest that Neanderthals matured earlier than modern humans. Ramirez Rozzi and Bermúdez de Castro (2004) found Neanderthal lateral enamel formation times to be about 15% shorter than in modern humans, and T. M. Smith et al. (2007, 2010) reported significantly earlier eruptions of Neanderthal M1 and M2 compared with both living and fossil *Homo sapiens*. These results point to a faster pace of dental, and possibly overall, development in Neanderthals, although other authors have argued that Neanderthal growth rates may fall within the modern human range, underscoring an ongoing debate (Cutler, 1975; Ponce de León et al., 2008, 2016). Thus, the timing of the evolution of extended adrenarche and middle childhood remains unresolved by morphological evidence.

Genetic analysis provides a complementary approach to fossil-based inferences of growth and life history, offering new insights into adrenarche and middle childhood development in modern humans and Neanderthals. In this study, we examine adrenal steroidogenic genes linked to adrenarche and DHEA/S production in *Homo sapiens* and Neanderthals, with particular attention to key biosynthetic loci (Figure 1). DHEA, often measured as circulating DHEAS, serves as a precursor for sex hormones such as testosterone and estrogen, and its elevation during adrenarche underlies many of the physical and cognitive changes characteristic of middle childhood.

**Figure 1.**
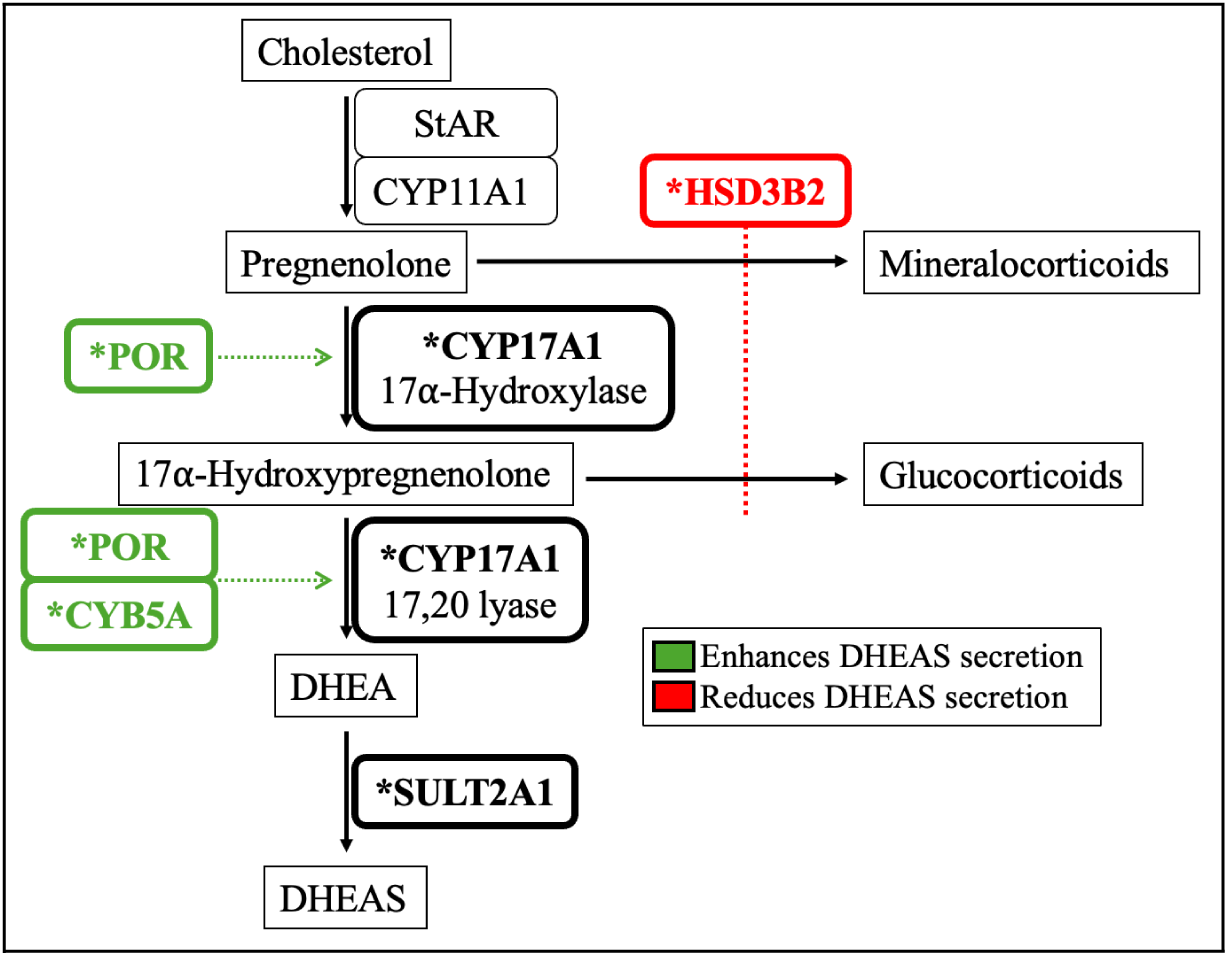
Steroidogenic pathway for the production of dehydroepiandrosterone sulfate (DHEAS) in the adrenal cortex. Asterisks indicate genes analyzed in this study, and abbreviations are explained in the text. This figure was modified from Rainey and Nakamura (2008).

The rise in adrenarche-defining DHEA/S results from increased steroidogenic activity in the ZR (Dhom, 1973; Nguyen & Conley, 2008). The first step in all steroid hormone biosynthesis is the conversion of cholesterol to pregnenolone, facilitated by the steroidogenic acute regulatory protein (StAR, encoded by *StAR*) and cytochrome P450scc (encoded by *CYP11A1*) (Figure 1, Figure S1) (Miller, 2002). DHEA/S is then synthesized from pregnenolone via the genes *CYP17A1* (cytochrome P450 17A1), *POR* (cytochrome P450 oxidoreductase), *CYB5A* (cytochrome b5 Type A), and *SULT2A1* (sulfotransferase Family 2A Member 1), while the expression of *HSD3B2* (3β-hydroxysteroid dehydrogenase type 2) acts a rate-limiting factor in DHEA/S formation (Figure 1) (Rainey & Nakamura, 2008). The onset of adrenarche begins with a regulatory shift within the ZR, particularly the increase in *CYP17A1* expression and the decrease of *HSD3B2* expression (Rainey & Nakamura, 2008; Rhéaume et al., 1991). In DHEA/S biosynthesis, the CYP17A1 enzyme functions as both 17α-hydroxylase and 17,20-lyase (Miller, 2002). POR works in both of these reactions as an obligate redox partner, and CYB5A allosterically enhances 17,20-lyase activity and can increase DHEA/S production up to 10-fold (Miller, 2002; Yoshimoto & Auchus, 2015). Finally, *SULT2A1* encodes the specific sulfotransferase enzyme that converts DHEA to DHEAS (Rainey & Nakamura, 2008). While Figure 1 illustrates the focused DHEA/S biosynthesis pathway examined in this study, these genes are situated within a broader steroid hormone biosynthesis network; a more complete pathway diagram is provided in Figure S1.

By analyzing genetic variation and introgression patterns between modern humans and Neanderthals for DHEA/S biosynthesis genes acting downstream of the universal conversion of cholesterol to pregnenolone - specifically *HSD3B2*, *CYP17A1, POR, CYB5A*, and *SULT2A1 -* we ask whether the prolonged adrenarche seen in modern humans represents a derived adaptation or reflects deeper hominin roots. We test two scenarios for its origin: (1) a shared origin, in which Neanderthals and modern humans exhibit similar genetic mechanisms and unimpeded introgression, implying a common ancestral form of adrenarche; and (2) a divergent origin, where functional variants and introgression barriers indicate that prolonged adrenarche evolved uniquely in modern humans after diverging from Neanderthals. Our comparative analysis helps reveal genetic factors underlying lineage variation in middle childhood, and to refine evolutionary origins of this critical life-history stage.

## Materials and Methods

All custom code used in this analysis is publicly available at https://github.com/nickhartman24/Adrenarche-Project. Specific software packages and versions used across all analytical workflows are detailed in Supplementary Table S1.

### Sourcing Genomes

Sequence data were obtained from the 1000 Genomes Project (1KG, Phase 3), representing diverse global populations. Variant call format (VCF) files were downloaded from the official FTP release (1000 Genomes Project Consortium, 2015). All human sequences are aligned to the hg19/GRCh37 reference genome to ensure consistency in coordinate mapping and downstream comparison. For analyses involving comparison to the chimpanzee genome, we retrieved the hg19.panTro6.synNet multiple alignment data from the UCSC Genome Browser, which provides pairwise sequence alignments between human and chimpanzee genomes (Kent et al., 2002). Neanderthal and Denisovan genomes were sourced from the Max Planck Institute for Evolutionary Anthropology’s genome projects portal (https://www.eva.mpg.de/genetics/genome-projects). These high-coverage genomes are aligned to the hg19/GRCh37 reference, which facilitates direct comparison with modern human data (Mafessoni et al., 2020; Meyer et al., 2012; Prüfer et al., 2014, 2017).

### Identifying Introgression

To identify Neanderthal introgression into modern human populations, we used a method independent of the assumption that African populations have no archaic ancestry. We utilized introgression maps generated by Steinrücken et al. (2018) via diCal-admix, a hidden Markov model (HMM) that incorporates explicit demographic history. We analyzed introgression maps for a total of 282 individuals (564 chromosomes) across three 1KG populations: Utah residents with Northern and Western European ancestry (CEU) (n=85, 2n=170), and amalgamated Han Chinese in Beijing (CHB) and Southern Han Chinese (CHS), termed CHBS (n=197, 2n=394) by Steinrücken et al. (2018). These maps partition the genomes into 500-bp windows across both chromosome copies, calculating the localized probability that each window derives from Neanderthal admixture. Windows were classified as definitively introgressed if their assigned Neanderthal probability met the original study’s validated threshold of 0.89. The final metric we utilized for later analysis was the introgression frequency at each locus, calculated as the proportion of definitively introgressed segments relative to the total number of chromosomes sampled per population (Table 2). We repeated the introgression map analysis for 5,000bp upstream and downstream of each gene to determine if introgressed haplotypes continued past the gene boundaries and into potential cis-regulatory regions (Table S5). The analysis of *POR* was truncated upstream by the start of the gene *TMEM120A*. We also conducted an introgression analysis using the Haplostrips software package to visualize haplotypes (Marnetto & Huerta-Sánchez, 2017). Full methods for haplotype analysis are described in Supplementary Methods, and visualized in Figure S2.

### Identifying Neanderthal-specific Variants

To identify Neanderthal-specific variants, we used chromosome-specific 1KG VCF files and high-coverage Vindija, Altai, and Chagyrskaya Neanderthal genomes from the Max Planck Institute, all aligned to the hg19/GRCh37 human reference genome. For each gene of interest, genomic intervals, including untranslated regions (UTRs), were defined using coordinates from the UCSC Genome Browser. Archaic VCFs were subsequently subset by gene and filtered to retain only single nucleotide variants (SNVs). Similarly, 1KG Yoruba (YRI) VCF was subset to the same gene-specific regions, and filtered to sites that overlapped SNV positions in the archaic files. After merging the YRI and archaic datasets, we utilized VCFtools to generate allele frequencies at each site. A site was classified as a Neanderthal-specific SNV if the allele was fixed across all Neanderthal genomes but entirely absent within the YRI cohort. This filtering strategy leverages the YRI as a reference group; because the Yoruba represent a deeply diverged, ancient modern human lineage with putatively minimal archaic admixture, utilizing them here effectively reduces confounding from shared ancestral variation and global introgression. To determine ancestral versus derived state for the modern human and Neanderthal alleles at each position, we used the UCSC Multiz Alignments of 100 Vertebrates (Blanchette et al., 2004). To assess whether Neanderthal-specific variants exist beyond the defined gene coordinates, we examined flanking upstream (5,000 bp) and downstream (5,000bp) regions for each of the five genes for putative cis-regulatory variation.

### Allele Frequencies of Archaic Variants

To calculate the population-level allele frequencies of the Neanderthal SNVs identified by our variant calling process, we adapted the open-source workflow developed by Wroblewski et al. (2023), which was originally designed to detect archaic SNVs within pharmacogenes (code available at https://github.com/kelsey-witt/archaic_pgx). Specifically, these scripts were tailored to target our five genes of interest. The allele frequency script identifies alleles present at low frequencies within 1KG African populations and the alternate alleles present in archaic genotypes for each target locus, which functioned as a confirmation of the variants identified in our earlier variant calling process. Subsequently, the comparative script produces a list of allele frequencies for all 1KG populations at these Neanderthal-derived SNVs (Figure 2, Table S2).

**Figure 2.**
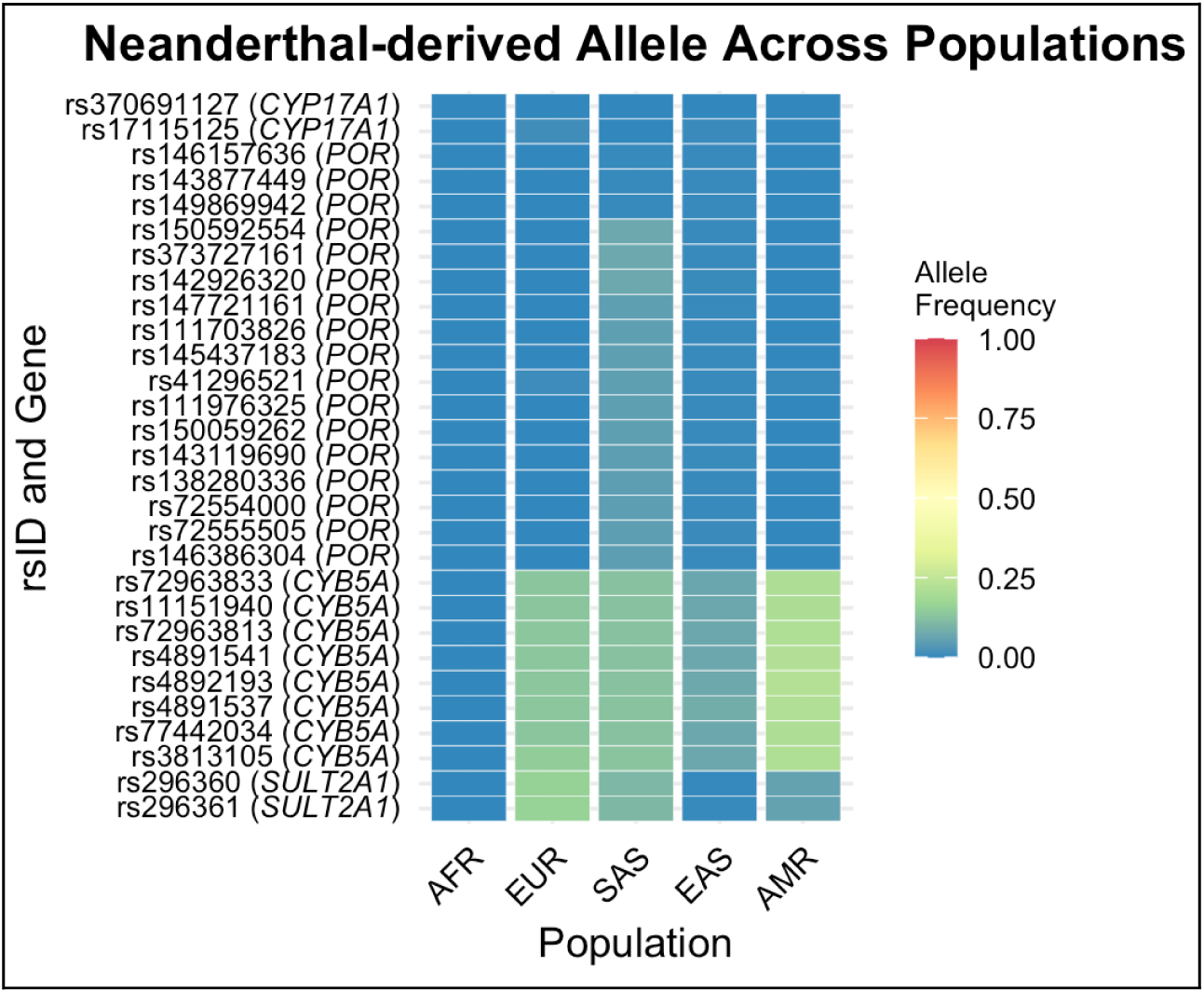
Allele frequency of Neanderthal variants in modern human superpopulations, as described by the 1000 Genomes Project. Superpopulations represented by AFR (Africans), EUR (Europeans), SAS (South Asians), EAS (East Asians), and AMR (Admixed Americans). Exact allele frequencies are available in Supplementary Table S3.

### Test for Positive Selection

We employed nf-selection, a Nextflow-based pipeline for detecting signatures of recent natural selection, to test for evidence of positive selection in introgressed loci (Miron-Toruno et al., 2025; GitHub: https://github.com/fernanda-miron/nf-selection). Analyses were restricted to the two populations exhibiting the highest allele frequencies for *CYB5A* and *SULT2A1*, and the highest allele frequency population for *POR*. The pipeline implements two complementary statistics: Population Branch Statistic (PBS) and integrated haplotype score (iHS). PBS quantifies population-specific allele frequency divergence across three populations; in our analyses, population 1 corresponded to the focal population, while East Asian (EAS) and African (AFR) populations from the 1000 Genomes Project served as comparison and outgroup populations, respectively. PBS values were computed genome-wide, and variants were considered candidates for positive selection if they fell within the top 1% of the empirical PBS distribution. Concurrently, iHS was calculated for each variant on chromosomes harboring genes of interest. The iHS statistic detects recent selection by measuring extended haplotype homozygosity within populations; variants were prioritized as candidates if they exhibited a p-value <0.01.

### Database search for function of variants

We conducted a systematic database search to identify relevant studies on the functional effects for all candidate Neanderthal-specific variants. Standardized reference SNP cluster IDs (rsID) were retrieved from the NCBI Single Nucleotide Polymorphism Database (dbSNP) (https://www.ncbi.nlm.nih.gov/snp/; Sherry et al., 2001). Tissue-specific expression and splicing consequences were searched using the GTEx Portal (Figure S3) (https://gtexportal.org; GTEx Consortium, 2020), while organism-level phenotypic associations were searched using the NHGRI-EBI GWAS Catalog (https://www.ebi.ac.uk/gwas; Welter et al., 2014). For a review of existing literature, we searched rsIDs and locus-specific keywords in LitVar 2.0 (https://www.ncbi.nlm.nih.gov/research/litvar2; Allot et al., 2023).

## Results

We identified 29 Neanderthal-derived single nucleotide variants (SNVs) across the adrenal steroidogenic genes examined (Table 1). 27 of these variants are located within the genic region, while 2 variants were identified upstream of *CYB5A* during our search for cis-regulatory variants. The modern human allele frequencies for all variants are illustrated in Figure 2 (allele frequencies available in Table S3). For each of these SNVs, we assessed the corresponding reference allele in *Pan troglodytes* (chimpanzee) and the Denisovan genome (Table 1), and we determined that modern humans maintain the ancestral state at each of these positions, while Neanderthals are derived for all SNVs (Table S4). Three genes were also identified as having introgressed Neanderthal haplotypes in some non-African modern human populations (Table 2).

**Table 1.** For each Neanderthal-like SNV identified, this table reports the hg19/GRCh37 position, rsID, and chimpanzee (Pan), modern human (Hum), Denisovan (Den), and Neanderthal (Nea) reference alleles. Nucleotides are color-coded: adenosine (blue), guanine (green), cytosine (yellow), and thymine (pink). Denisovan is heterozygous at 6 positions in *POR*.

| Gene | Position | rsID | Pan | Hum | Den | Nea | Location |
| --- | --- | --- | --- | --- | --- | --- | --- |
| *CYP17A1* | 10:104597159 | rs370691127 | T | T | T | A | 5' UTR exon |
| *CYP17A1* | 10:104597132 | rs17115125 | T | C | C | T | 5' UTR exon |
| *POR* | 7:75547674 | rs146157636 | A | A | A/T | T | intron |
| *POR* | 7:75548020 | rs143877449 | G | G | G/C | C | intron |
| *POR* | 7:75552167 | rs149869942 | T | T | T/G | G | intron |
| *POR* | 7:75565514 | rs150592554 | C | C | C/T | T | intron |
| *POR* | 7:75569375 | rs373727161 | C | C | C/T | T | intron |
| *POR* | 7:75579560 | rs142926320 | C | C | C/T | T | intron |
| *POR* | 7:75595927 | rs147721161 | G | G | G | T | intron |
| *POR* | 7:75597522 | rs111703826 | G | G | G | A | intron |
| *POR* | 7:75597561 | rs145437183 | G | G | G | C | intron |
| *POR* | 7:75598855 | rs41296521 | C | C | C | T | intron |
| *POR* | 7:75598872 | rs111976325 | G | G | G | T | intron |
| *POR* | 7:75599662 | rs150059262 | G | C | C | T | intron |
| *POR* | 7:75607167 | rs143119690 | G | G | G | A | intron |
| *POR* | 7:75607681 | rs138280336 | A | A | A | G | intron |
| *POR* | 7:75609144 | rs72554000 | G | G | A | A | intron |
| *POR* | 7:75609848 | rs72555505 | A | G | G | A | intron |
| *POR* | 7:75611935 | rs146386304 | C | C | C | A | intron |
| *CYB5A* | 18:71958555 | rs72963833 | G | G | G | A | intron |
| *CYB5A* | 18:71955452 | rs11151940 | C | C | C | A | intron |
| *CYB5A* | 18:71948446 | rs72963813 | G | G | A | A | intron |
| *CYB5A* | 18:71947422 | rs4891541 | C | C | C | T | intron |
| *CYB5A* | 18:71943304 | rs4892193 | T | T | T | C | intron |
| *CYB5A* | 18:71936427 | rs4891537 | C | C | C | T | intron |
| *CYB5A* | 18:71959324 | rs77442034 | G | G | G | A | upstream |
| *CYB5A* | 18:71960202 | rs3813105 | C | C | C | G | upstream |
| *SULT2A1* | 19:48388658 | rs296360 | T | T | T | C | intron |
| *SULT2A1* | 19:48389363 | rs296361 | G | G | G | A | intron |

### HSD3B2

For *HSD3B2*, our analyses showed no evidence of archaic introgression, and no Neanderthal-derived SNVs were detected within *HSD3B2*.

### CYP17A1

Our introgression analysis of *CYP17A1* yielded no signal of introgression at this locus in any modern human population. Through variant calling, two Neanderthal-derived SNVs within the 5′ untranslated region (UTR) exons of CYP17A1 were identified. One variant, rs17115125, is restricted to East Asian (0.68%) and European (1.2%) superpopulations (Figure 2). Given the low frequency of this allele and the lack of a surrounding introgressed haplotype background, this site likely represents an instance of recurrent mutation rather than true archaic introgression. An automatic clinical scoring system applied by Illumina Laboratory Services in 2018 classified the rs17115125 T allele as a variant of uncertain significance for steroid 17-alpha-monooxygenase (CYP17A1/cytochrome P450c17) deficiency in adults (Accession: RCV000357830.5; NCBI, 2018). The second Neanderthal-derived SNV, rs370691127, was not observed in any 1KG individuals (Figure 2), and no clinical reports were identified for this site.

### POR

We identified 17 Neanderthal-derived variants in *POR*, which are all intronic. All variants are present in SAS (0.4 - 7.2 %) and EAS populations (0.5 - 0.7%), and are broadly absent in all other modern human populations (Figure 2, Table S3). Although the introgression maps do not include South Asian populations, one chromosome in the CHBS sample group was classified as likely introgressed (Table 2). In the Haplostrips haplotype analyses, several South Asian individual haplotypes showed greater similarity to Neanderthal haplotypes, although none formed a close match to the Neanderthal sequence (Figure S2). Despite this, the fact that all identified Neanderthal-matching variants are present in modern humans at moderate frequency suggests that the gene may retain an introgressed archaic signal. Because the introgression maps are limited to CEU and CHBS populations, we were unable to determine whether the Neanderthal haplotype observed in South Asians extends into the flanking regions of the gene. The scan for selection using nf-selection did not identify any variants meeting the significance thresholds for PBS or iHS for any population analyzed (Table S6). No adrenal associations were identified in the database search for any of the *POR* Neanderthal-derived variants.

**Table 2.** Introgression frequencies at each of the five DHEA/S biosynthesis genes examined, derived from Neanderthal introgression maps generated by Steinrücken et al. (2018) via diCal-admix. Results are shown separately for Central European (CEU) and aggregated East Asian (CHBS) populations.

| Gene | Chr | Start Pos | End Pos | Population | # of individuals | Total chr copies | Sum introgressed chr copies | Introgression Frequency |
| --- | --- | --- | --- | --- | --- | --- | --- | --- |
| HSD3B2 | 1 | 119957773 | 119965657 | CEU | 85 | 170 | 0 | 0.0000000 |
|  |  |  |  | CHBS | 197 | 394 | 0 | 0.0000000 |
| CYP17A1 | 10 | 104590288 | 104597170 | CEU | 85 | 170 | 0 | 0.0000000 |
|  |  |  |  | CHBS | 197 | 394 | 0 | 0.0000000 |
| POR | 7 | 75544475 | 75616173 | CEU | 85 | 170 | 0 | 0.0000000 |
|  |  |  |  | CHBS | 197 | 394 | 1 | 0.002538071 |
| CYB5A | 18 | 71918081 | 71959198 | CEU | 85 | 170 | 30 | 0.1764706 |
|  |  |  |  | CHBS | 197 | 394 | 41 | 0.104060914 |
| SULT2A1 | 19 | 48373724 | 48389572 | CEU | 85 | 170 | 31 | 0.1823529 |
|  |  |  |  | CHBS | 197 | 394 | 1 | 0.002538071 |

### CYB5A

Introgression maps support introgression at *CYB5A*, with introgression frequencies differing between CEU (17.6%) and aggregated CHBS (10.4%) populations (Table 2). The introgressed region also continued across the flanking regions of *CYB5A* (Table S5). We identified six intronic Neanderthal-derived variants in *CYB5A*, along with two upstream variants (Table 1). All eight variants are present in non-African superpopulations at varying allele frequencies (Figure 2, Table S3). In East Asian populations, variant frequencies ranged from 6.84 - 8.34%; in South Asians, variants were present in 12.5-13.2% of individuals. European populations carried these variants at 13.4 - 14.8%, and Admixed American populations at 20.9 - 22.3%. Despite these distributions, no variants met the predefined significance thresholds for PBS or iHS in any population analyzed using nf-selection (Table S6). No adrenal associations were identified in the database search for any of the genic *CYB5A* Neanderthal-derived variants. However, the two Neanderthal-derived variants (rs77442034, rs3813105) located upstream of the gene were identified as splicing quantitative trait locus (sQTL) in GTEx, associated with altered splicing of the gene in the adrenal gland, consistent with tissue-specific cis-regulatory effects.

### SULT2A1

For *SULT2A1*, introgression analyses show signals of introgression at this locus, with marked differences between CEU (18.2%) and aggregated CHBS (0.25%) populations (Table 2). The introgressed region also continued into the *SULT2A1* flanking regions (Table S5).Within *SULT2A1*, we identified two intronic Neanderthal-derived variants. Both variants are present in modern humans through introgression, with allele frequencies between 15.9 - 16.1% in European populations and 10.4-10.5% in South Asian populations (Figure 2, Table S3). For both variants, the Neanderthal-derived allele was present in 5.9% of Admixed American populations and in less than 0.5% of East Asian populations. No variants met the predefined significance thresholds for PBS or iHS in any population analyzed using nf-selection (Table S6). Database searches identified clinical associations for each variant. The rs296361 introgressed A allele has been associated with decreased levels of a steroid metabolite, 5α-pregnan-3β,20α-diol disulfate (Hysi et al., 2022). Additionally, For rs296360, one adult cohort study reported that the introgressed C allele was associated with a 0.1427-unit decrease in DHEAS levels (Pott et al., 2019). According to GTEx, both rs296361 and rs296360 are adrenal gland expression quantitative trait loci (eQTLs) for *SULT2A1*. Heterozygotes carrying the human and Neanderthal alleles show significantly reduced *SULT2A1* RNA expression in adrenal tissue, suggesting that these variants may influence adrenal expression of the sulfotransferase.

## Discussion

Compared to other apes, modern humans exhibit a uniquely prolonged adrenarche, which is thought to have evolved to promote and extend cognitive and social development (Bogin, 1997). An important question in life history theory is when this uniquely prolonged adrenarche first appeared in our evolutionary history: was it exclusive to *Homo sapiens*, or present in other hominins such as Neanderthals? Based on fossil evidence, there is ongoing debate over whether Neanderthals developed faster than modern humans or if they fall within the modern human range (Cutler, 1975; Ponce de León et al., 2008, 2016; Ramirez-Rozzi & Bermúdez de Castro, 2004; T. M. Smith et al. 2007, 2010). To address this gap, we use a comparative genetic approach focused specifically on genes in the DHEA/S biosynthesis pathway, as DHEA is the primary hormone driving changes during adrenarche, and variation in DHEA/S levels directly corresponds to differences in adrenarche duration among primates (Bernstein et al., 2012; Nguyen & Conley, 2008).

Our genetic analysis identified 29 Neanderthal-derived single nucleotide variants (SNVs) in non-coding regions across the adrenal steroidogenic genes examined. We also looked for evidence of introgression into modern humans, as genes with identical function should be interchangeable between the two species, and natural selection would not have removed them from the modern human gene pool (or raise their frequency in case of adaptation). Rather than a uniform pattern, introgression across the DHEA/S biosynthesis pathway appears to be constrained by each gene’s functional role. Central catalytic genes such as *HSD3B2* and *CYP17A1* show no evidence of introgression, whereas accessory redox partners (*POR, CYB5A*) and the sulfating enzyme (*SULT2A1*) carry introgressed Neanderthal haplotypes at low-to-moderate frequencies in non-African populations.

Because the central biosynthetic genes *HSD3B2* and *CYP17A1* function across multiple steroid hormone biosynthetic pathways (Figure S1), introgression at these loci may be constrained by their pleiotropic roles. Because *HSD3B2* is essential for synthesizing all major steroid hormone classes in the adrenal glands and gonads, broad protein-coding sequence conservation across primates is the most parsimonious explanation for lack of variation at this gene (Bernstein et al., 2012; Simard et al., 2005). In contrast, we identified distinct sequence differences between modern humans and Neanderthals for *CYP17A1* that were not introgressed - with rs17115125 more likely reflecting recurrent mutation. Because CYP17A1 functions in both DHEA/S biosynthesis and other critical steroid pathways (Figure S1), introgressed archaic variants may introduce functional incompatibilities, leading to negative selection against Neanderthal alleles at this locus.

The genes *POR, CYB5A*, and *SULT2A1*, while active across diverse biological contexts, execute tissue-dependent regulatory roles within the adrenal gland (Figures S1, S3). Specifically, *POR* and *CYB5A* act as redox partners and allosteric regulators for steroidogenesis, while *SULT2A1* functions as a downstream sulfotransferase (Miller, 2002). Unlike the key catalytic genes, these accessory functions may tolerate variation in Neanderthal-derived alleles without widespread negative effects. The introgression of Neanderthal-derived variants across *POR, CYB5A*, and *SULT2A1* suggests functional compatibility between modern and archaic lineages, with these variants either preserving core enzymatic function or exerting regulatory effects too subtle to be physiologically disruptive.

All Neanderthal-specific variants identified in this study fall in non-coding regions, consistent with the well-established pattern that phenotypic differences between modern humans and Neanderthals are often driven by gene expression differences rather than protein-coding changes (Colbran et al., 2019; McCoy et al., 2017). Three variants are worth highlighting. The *SULT2A1* variant rs296361 has been associated with decreased levels of 5α-pregnan-3β,20α-diol disulfate, consistent with altered *SULT2A1*-dependent steroid sulfation, and rs296360 has been associated with decreased DHEAS levels in adult women (Hysi et al., 2022; Lightning et al., 2021; Pott et al., 2019). Both variants are eQTLs, with Neanderthal allele carriers showing reduced *SULT2A1* expression. The *CYP17A1* variant rs17115125 has been associated with CYP17A1 enzyme deficiency in adults, suggesting a potential shift in steroidogenic efficiency (Accession: RCV000357830.5; NCBI, 2018). Together, this raises the possibility that Neanderthals carrying these alleles may have had modestly lower DHEA/S levels, though effects during adrenarche specifically remain unknown.

When compared with chimpanzee, Denisovan, and other primate alignments to determine ancestral character states and identify lineage-specific changes (Table 1), we found that modern humans maintain the ancestral allele at all 29 variant positions (Table S4). Alternatively, the Neanderthals display an array of lineage-specific derived variants. Because modern humans retained the ancestral allele across the variant positions, we find no evidence for a sweep of novel, human-specific variants driving our extended adrenarche compared to other hominins.

Returning to the central question of this study, our results offer insights into the origin and timing of extended human adrenarche. Because all identified Neanderthal-derived variants are located within non-coding regions, any physiological differences in adrenarche between these species would likely stem from differential gene expression rather than structural divergence in the enzymes themselves. Consistent with this, three of these Neanderthal-derived variants are known to be associated with modestly reduced enzyme activity or lower DHEA/S production in modern humans. This could suggest a shorter or earlier adrenarche in Neanderthals compared to humans, an interpretation that aligns with fossil evidence of accelerated Neanderthal development. The broader introgression pattern showed that while accessory genes exhibit signs of archaic introgression in modern human populations, the core biosynthetic genes do not, reinforcing that genes with narrower pleiotropic effects are more flexible to genetic variation. Finally, our analysis of cis-regulatory regions revealed Neanderthal-derived variants in the flanking sequences of only one gene, *CYB5A*, and the introgression of these variants into modern humans suggests functional compatibility with the modern human genomic background. The absence of cis-regulatory introgression at the remaining loci suggests that expression differences elsewhere are more likely attributable to trans-regulatory mechanisms or epigenetic variation.

Several important limitations must be considered when interpreting these findings. Genetic factors, while influential, are not deterministic - the timing of adrenarche is also substantially affected by environmental factors such as nutrition, birth weight, obesity, and stress, as well as by epigenetic modifications, all of which can shape both the onset and extent of adrenarche in complex ways (Rosenfield, 2021).

## Conclusion

By investigating archaic introgression and genetic variation of the DHEA/S biosynthesis pathway, this study provides novel genetic insight into the evolutionary origins of extended adrenarche and middle childhood. Through a comparative analysis of five key steroidogenic genes between Neanderthals and globally diverse modern humans, we find a pattern in which central biosynthetic genes (*HSD3B2, CYP17A1*) show no evidence of Neanderthal introgression, while accessory genes (*POR, CYB5A, SULT2A1*) tolerate introgressed Neanderthal haplotypes at low-to-moderate frequencies. The sequence conservation of HSD3B2 across lineages, together with the successful introgression of variants at the three accessory loci, points to functional compatibility between Neanderthal and modern human DHEA/S biosynthesis genes. Furthermore, the absence of Neanderthal-derived variants in protein-coding exons across all five genes points to regulatory divergence as the likely basis of any interspecific variation in adrenarche. Three introgressed variants are associated with mildly reduced enzyme expression or DHEA/S levels, suggesting that Neanderthal adrenarche may have been slightly reduced relative to the modern human pattern, though not fundamentally different. Crucially, because modern humans preserve the ancestral ape allele at all key loci, modern human’s extended adrenarche does not reflect a sweep of novel, human-specific mutations. Ultimately, our results indicate that extended adrenarche is not a uniquely modern human innovation; but at least the genetic underpinnings of the DHEA/S pathway are shared between modern humans and Neanderthals.

Future research can expand upon these findings by moving beyond the DHEA/S biosynthesis pathway to encompass distant regulatory regions and the potential impacts of epistasis. Because the steroidogenic genes analyzed here are positioned within highly integrated steroid biosynthesis and metabolic networks, future studies must evaluate how epistatic interactions between these introgressed variants and the broader genetic background shape variation in DHEA/S production.

