## Supplementary Materials for "Shared genetic foundation of the DHEAS pathway indicate similar extended childhoods in Neanderthals and modern humans"

#### Table of Contents:

Supplementary Methods

- Haplostrips methods
- Table S1: Software packages

Supplementary Tables and Figures

- Figure S1: Steroid hormone biosynthesis pathway
- Figure S2: Haplotype genetic distance
- Figure S3: Bulk-tissue gene expression
- Table S2: Archaic-derived variants and allele frequencies
- Table S3: Neanderthal-derived variants
- Table S4: Variant ancestral/derived allele states
- Table S5: Introgression frequencies
- Table S6: nf-selection results

References

### Supplementary Methods

*Visualizing haplotypes using Haplostrips*

To assess introgression patterns, we conducted haplotype analysis using the software Haplostrips (Marnetto & Huerta-Sánchez, 2017). We analyzed diploid human samples from 1KG (n = 1,951, 2n = 3,902) alongside three Neanderthal genomes, designating the Vindija Neanderthal as the reference for haplotype comparisons. Gene coordinates were based on the hg19 assembly and included untranslated regions (UTRs). We applied a minor allele frequency threshold of 0.001 to retain rare variants. Haplostrips generated clustered haplotype plots for each genomic region, displaying haplotype relationships between the Vindija Neanderthal and all 1KG individuals. The software also calculated pairwise genetic distance between each 1KG individual and the Vindija Neanderthal reference. We visualized these distance distributions using ridgeline plots separated by population (Figure S2).

**Table S1.** Software packages, versions, and purposes across all analytical workflows. For each tool, the relevant version(s) and primary purpose within the workflow are indicated.

| Software | Version | Method | Purpose in Workflow |
|---|---|---|---|
| Haplostrips | 1.3 | Haplostrips | Haplotype clustering and sorting |
| pandas | 0.24.2 | Haplostrips | Haplostrips preferred version |
| | 3.0.0 | Introgression maps | Data table management |
| python | 2.7.18 | Haplostrips | Haplostrips preferred version |
| | 3.11.14 | Variant identification | Variant frequency analysis |
| bcftools | 1.20 | Variant identification | VCF filtering |
| VCFtools | 0.1.14 | Variant identification | Allele frequency calculation |
| | 0.1.16 | nf-selection | $F_{st}$ calculation |
| Nextflow | 21.0.6 | nf-selection | Pipeline orchestration |
| R | 4.2.2 | nf-selection | PBS, iHS calculation |
| r-ggplot2 | 3.5.1 | nf-selection | PBS calculation; plots development |
| r-tidyr | 1.3.1 | nf-selection | preprocessing and PBS calculation; data frame manipulation |
| r-cowplot | 1.1.3 | nf-selection | PBS calculation; plots development |
| r-rehh | 3.2.3 | nf-selection | iHS computation |
| r-vroom | 1.6.5 | nf-selection | Data processing |
| r-circlize | 0.4.16 | nf-selection | Results visualization |

### Supplementary Tables and Figures

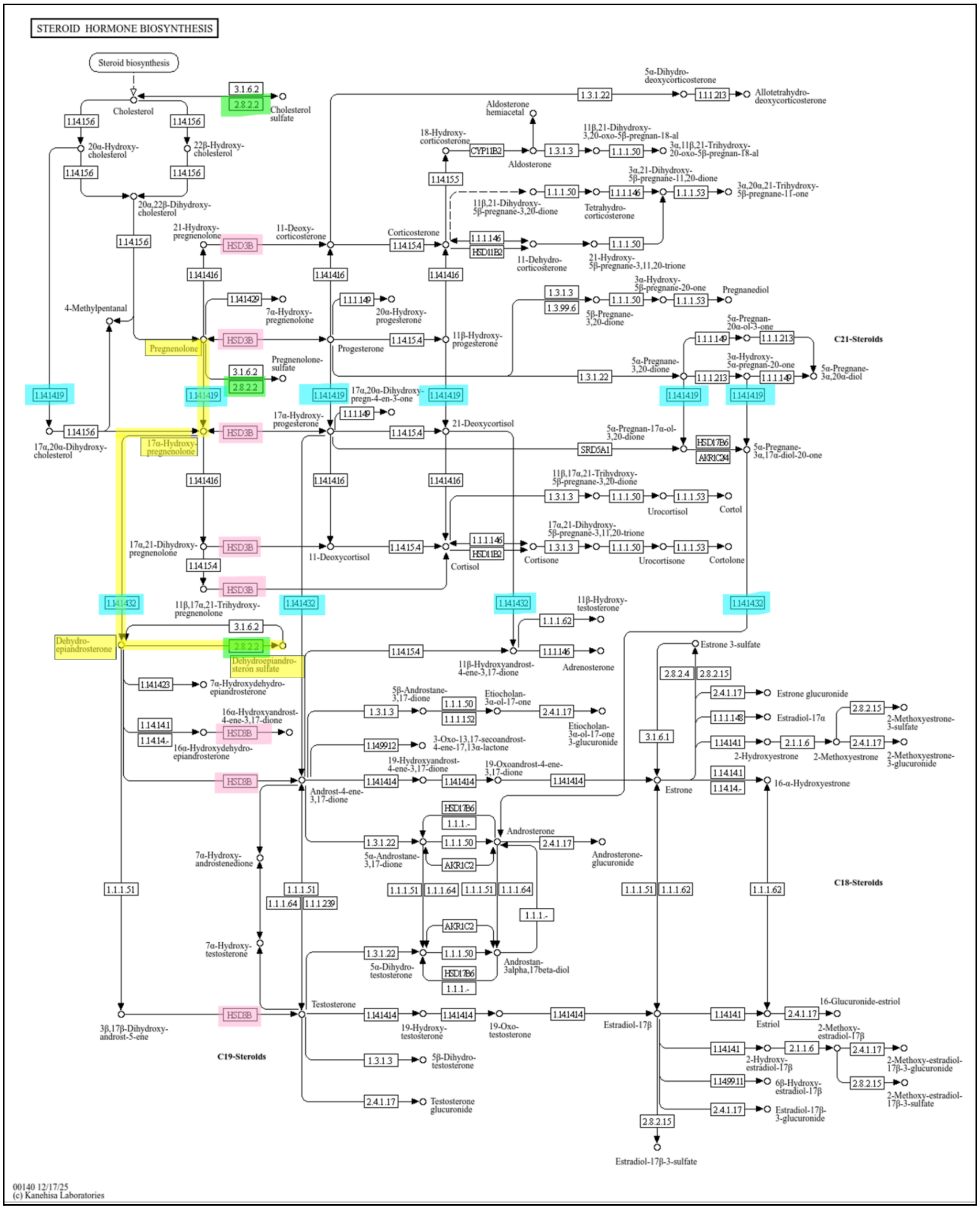


**Figure S1.** Steroid hormone biosynthesis pathway highlighting genes relevant to DHEA/S production. Yellow indicates the DHEA/S biosynthesis path; pink indicates HSD3B action (*HSD3B2* in the adrenal gland); green indicates *SULT2A1* enzyme action; blue indicates

*CYP17A1* enzyme action. *POR* and *CYB5A,* as redox partners, were not included in the diagram. Reproduced from the KEGG Pathway Database (Kanehisa et al., 2023).

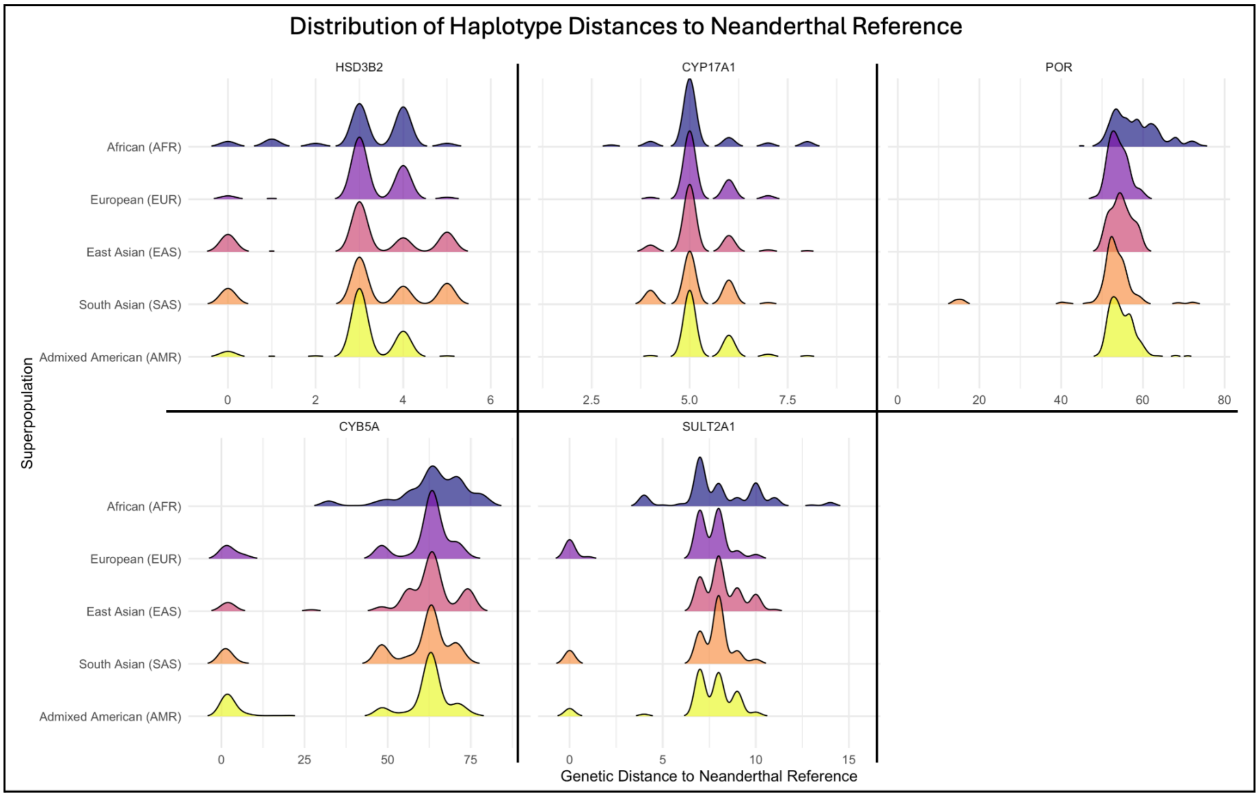


**Figure S2.** Distribution of pairwise genetic distances, as determined by Haplostrips, to the Vindija Neanderthal reference genome across five DHEA/S biosynthesis genes. Each ridgeline represents the distribution of genetic distances from the Vindija Neanderthal reference for a given superpopulation.

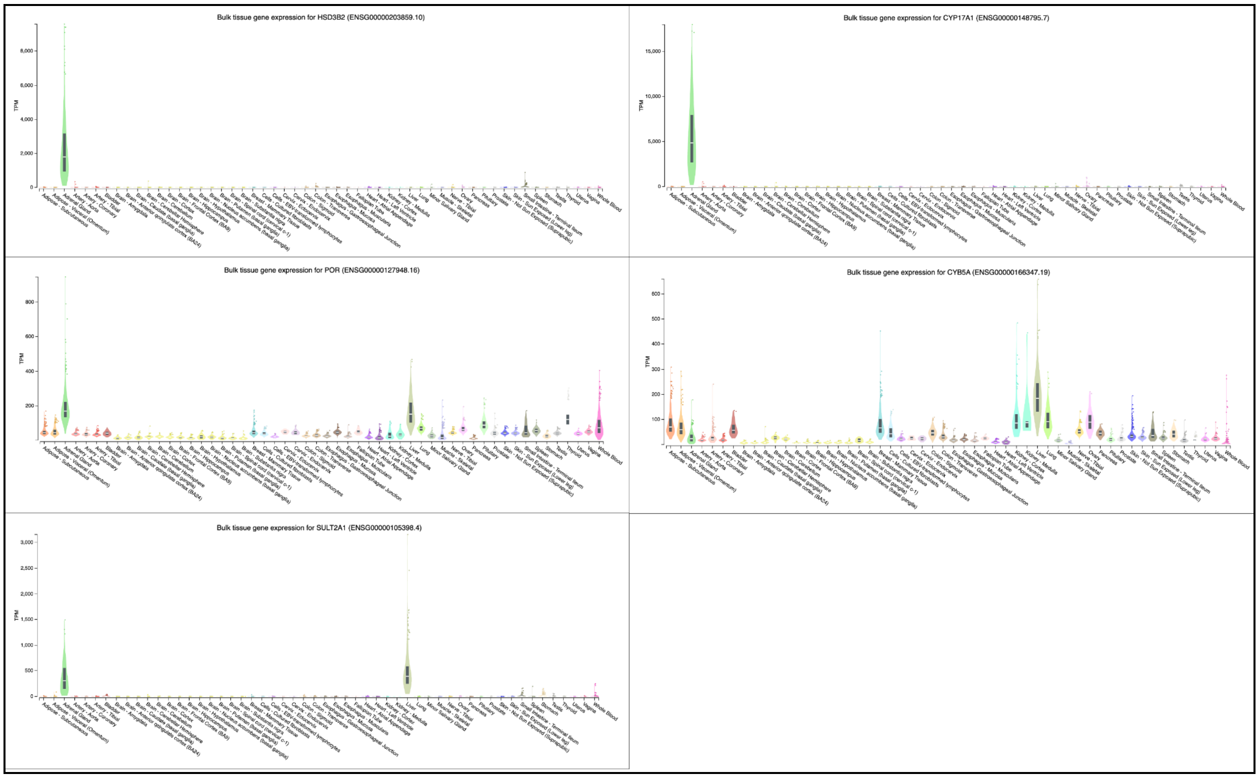


**Figure S3.** Bulk tissue gene expression for each of the five DHEA/S biosynthesis genes investigated in this study. Expression values (TPM) are shown across 54 human tissues. Data retrieved from the GTEx Portal (v8;GTEx Consortium, 2020).

| Gene | Chr | Position | rsID | Ref | Alt | Den | Altai | Chag | Vind | Arch _Alt | AFR | CHB | JPT | CHS | CDX | KHV | CEU | TSI | FIN | GBR | IBS | MXL | PUR | CLM | PEL | GIH | PJL | BEB | STU | ITU |
|---|---|---|---|---|---|---|---|---|---|---|---|---|---|---|---|---|---|---|---|---|---|---|---|---|---|---|---|---|---|---|

**extended_archaic_mp_nonafr_snp_freqs**

**Table S2.** All archaic-derived variants generated by the archaic-pgx variant calling pipeline across five DHEA/S biosynthesis genes, including variants within the 5kb flanking regions of each gene. For each site, the table reports the hg19/GRCh37 chromosomal position, rsID, reference and alternate alleles, Denisovan and Neanderthal (Altai, Chagyrskaya, Vindija) genotypes, and allele frequencies across 1000 Genomes Project populations. Rows highlighted in gray indicate variants meeting the filtering criteria for classification as Neanderthal-derived: allele frequency $\leq 0.001$ in African populations, and homozygous alternate genotype (1/1) in all three Neanderthal individuals. Population abbreviations follow 1000 Genomes Project conventions.

| Gene | Chr | hg19 position | rs number | ref | Nea. Alt | AFR | SAS | EAS | EUR | AMR |
|---|---|---|---|---|---|---|---|---|---|---|
| *CYP17A1* | 10 | 104597132 | rs17115125 | C | T | 0 | 0.000 | 0.007 | 0.012 | 0.000 |
| *CYP17A1* | 10 | 104597159 | rs370691127 | T | C | 0.001 | 0.000 | 0 | 0.000 | 0.000 |
| *POR* | 7 | 75547674 | rs146157636 | A | T | 0 | 0.004 | 0.005 | 0 | 0 |
| *POR* | 7 | 75548020 | rs143877449 | G | C | 0 | 0.004 | 0.005 | 0 | 0 |
| *POR* | 7 | 75552167 | rs149869942 | T | G | 0 | 0.004 | 0.005 | 0 | 0 |
| *POR* | 7 | 75565514 | rs150592554 | C | T | 0 | 0.071 | 0.005 | 0 | 0 |
| *POR* | 7 | 75569375 | rs373727161 | C | T | 0 | 0.071 | 0.005 | 0 | 0 |
| *POR* | 7 | 75579560 | rs142926320 | C | T | 0 | 0.072 | 0.006 | 0 | 0 |
| *POR* | 7 | 75595927 | rs147721161 | G | T | 0 | 0.051 | 0.005 | 0 | 0 |
| *POR* | 7 | 75597522 | rs111703826 | G | A | 0 | 0.051 | 0.005 | 0 | 0 |
| *POR* | 7 | 75597561 | rs145437183 | G | C | 0 | 0.051 | 0.005 | 0 | 0 |
| *POR* | 7 | 75598855 | rs41296521 | C | T | 0 | 0.051 | 0.005 | 0.01 | 0.005 |
| *POR* | 7 | 75598872 | rs111976325 | G | T | 0 | 0.051 | 0.005 | 0 | 0 |
| *POR* | 7 | 75599662 | rs150059262 | C | T | 0 | 0.051 | 0.005 | 0 | 0 |
| *POR* | 7 | 75607167 | rs143119690 | G | A | 0 | 0.049 | 0.006 | 0 | 0 |
| *POR* | 7 | 75607681 | rs138280336 | A | G | 0 | 0.049 | 0.005 | 0 | 0 |
| *POR* | 7 | 75609144 | rs72554000 | G | A | 0 | 0.049 | 0.005 | 0 | 0 |
| *POR* | 7 | 75609848 | rs72555505 | G | A | 0.001 | 0.05 | 0.007 | 0 | 0 |
| *POR* | 7 | 75611935 | rs146386304 | C | A | 0 | 0.049 | 0.005 | 0 | 0 |
| *CYB5A* | 18 | 71936427 | rs4891537 | C | T | 0 | 0.126 | 0.083 | 0.135 | 0.217 |
| *CYB5A* | 18 | 71943304 | rs4892193 | T | C | 0 | 0.125 | 0.073 | 0.135 | 0.223 |
| *CYB5A* | 18 | 71947422 | rs4891541 | C | T | 0 | 0.125 | 0.071 | 0.136 | 0.219 |
| *CYB5A* | 18 | 71948446 | rs72963813 | G | A | 0 | 0.125 | 0.071 | 0.136 | 0.212 |
| *CYB5A* | 18 | 71955452 | rs11151940 | C | A | 0 | 0.125 | 0.07 | 0.134 | 0.209 |

| | | | | | | | | | | |
|---|---|---|---|---|---|---|---|---|---|---|
| *CYB5A* | 18 | 71958555 | rs72963833 | G | A | 0 | 0.125 | 0.068 | 0.134 | 0.210 |
| *CYB5A* | 18 | 71959324 | rs77442034 | G | A | 0 | 0.1252 | 0.0696 | 0.134 | 0.21 |
| *CYB5A* | 18 | 71960202 | rs3813105 | C | G | 0 | 0.132 | 0.0696 | 0.148 | 0.212 |
| *SULT2A1* | 19 | 48388658 | rs296360 | T | C | 0.001 | 0.104 | 0.003 | 0.159 | 0.059 |
| *SULT2A1* | 19 | 48389363 | rs296361 | G | A | 0.001 | 0.105 | 0.004 | 0.161 | 0.059 |

**Table S3.** Neanderthal-derived allele frequencies for the 29 identified Neanderthal-derived variants, averaged across five 1000 Genomes Project superpopulations (AFR, SAS, EAS, EUR, AMR).

| Gene | # variants | region | Chimpanzee | Modern Human | Denisovan | Neanderthal |
|---|---|---|---|---|---|---|
| *CYP17A1* | 1 | 5' UTR | A | A | A | D |
| *CYP17A1* | 1 | 5' UTR | D | A | A | D |
| *POR* | 8 | intron | A | A | A | D |
| *POR* | 6 | intron | A | A | A/D | D |
| *POR* | 2 | intron | D | A | A | D |
| *POR* | 1 | intron | A | A | D | D |
| *CYB5A* | 5 | intron | A | A | A | D |
| *CYB5A* | 1 | intron | A | A | D | D |
| *CYB5A* | 2 | upstream | A | A | A | D |
| *SULT2A1* | 2 | intron | A | A | A | D |

**Table S4:** Summarized ancestral/derived allele states, expanded on in-text Table 1. Ancestral (A), Derived (D), heterozygous (A/D). UCSC Multiz Alignments of 100 Vertebrates was used to determine ancestral vs derived state (Blanchette et al., 2004).

| Region | Chr | Start Pos | End Pos | Population | # of individuals | Total chr copies | Sum introgressed chr copies | Introgression Frequency |
|---|---|---|---|---|---|---|---|---|
| Upstream | 1 | 119952773 | 119957773 | CEU | 85 | 170 | 0 | 0.0000000 |
| *HSD3B2* | | | | CHBS | 197 | 394 | 0 | 0.0000000 |
| Downstream | 1 | 119965657 | 119970657 | CEU | 85 | 170 | 0 | 0.0000000 |
| *HSD3B2* | | | | CHBS | 197 | 394 | 0 | 0.0000000 |
| Upstream | 10 | 104585288 | 104590288 | CEU | 85 | 170 | 0 | 0.0000000 |
| *CYP17A1* | | | | CHBS | 197 | 394 | 0 | 0.0000000 |
| Downstream | 10 | 104597170 | 104602170 | CEU | 85 | 170 | 0 | 0.0000000 |
| *CYP17A1* | | | | CHBS | 197 | 394 | 0 | 0.0000000 |
| Upstream | 7 | 75539473 | 75544473 | CEU | 85 | 170 | 0 | 0.0000000 |
| *POR* | | | | CHBS | 197 | 394 | 0 | 0.0000000 |
| Upstream | 18 | 71959198 | 71964198 | CEU | 85 | 170 | 22 | 0.1294118 |
| *CYB5A* | | | | CHBS | 197 | 394 | 31 | 0.07868 |
| Downstream | 18 | 71918081 | 71913081 | CEU | 85 | 170 | 28 | 0.164706 |
| *CYB5A* | | | | CHBS | 197 | 394 | 36 | 0.09137 |
| Upstream | 19 | 48389572 | 48394572 | CEU | 85 | 170 | 30 | 0.1764706 |
| *SULT2A1* | | | | CHBS | 197 | 394 | 1 | 0.005882353 |
| Downstream | 19 | 48368724 | 48373724 | CEU | 85 | 170 | 26 | 0.1529412 |
| *SULT2A1* | | | | CHBS | 197 | 394 | 1 | 0.005882353 |

**Table S5.** Neanderthal introgression frequencies in 5,000 base pair flanking regions of the five DHEA/S biosynthesis genes examined, derived from Neanderthal introgression maps generated by Steinrücken et al. (2018) via diCal-admix. The downstream flanking region of *POR* was excluded because the gene is immediately adjacent to *TMEM120A*. Results are shown separately for Central European (CEU) and aggregated East Asian (CHBS) populations.

| Gene | Population | rsID | PBS value | iHS value | iHS p-value |
|---|---|---|---|---|---|
| *POR* | SAS | rs146157636 | 0 | N/A | N/A |
| *POR* | SAS | rs143877449 | 0 | N/A | N/A |

| | | | | | |
|---|---|---|---|---|---|
| *POR* | SAS | rs149869942 | 0 | N/A | N/A |
| *POR* | SAS | rs150592554 | 0.0704416 | 0.33805415 | 0.73532238 |
| *POR* | SAS | rs373727161 | 0.0704416 | 0.26599261 | 0.7902449 |
| *POR* | SAS | rs142926320 | 0.0697737 | 0.12172668 | 0.90311548 |
| *POR* | SAS | rs147721161 | 0.0470902 | 2.04880315 | 0.04048137 |
| *POR* | SAS | rs111703826 | 0.0470902 | 1.72201746 | 0.08506636 |
| *POR* | SAS | rs145437183 | 0.0470902 | 1.82520514 | 0.06797008 |
| *POR* | SAS | rs41296521 | 0.0470902 | 1.38100946 | 0.16727605 |
| *POR* | SAS | rs111976325 | 0.0470902 | 1.37923763 | 0.1678215 |
| *POR* | SAS | rs150059262 | 0.0470902 | 1.43614908 | 0.15095992 |
| *POR* | SAS | rs143119690 | 0.0430467 | N/A | N/A |
| *POR* | SAS | rs138280336 | 0.0447785 | N/A | N/A |
| *POR* | SAS | rs72554000 | 0.0447785 | N/A | N/A |
| *POR* | SAS | rs72555505 | 0.0422687 | 0.97125291 | 0.331422351 |
| *POR* | SAS | rs146386304 | 0.0447785 | N/A | N/A |
| *CYB5A* | AMR | rs72963833 | 0.1512925 | 1.243158 | 0.213809 |
| *CYB5A* | AMR | rs11151940 | 0.1480251 | 0.7709722 | 0.4407234 |
| *CYB5A* | AMR | rs72963813 | 0.1512925 | 0.2794105 | 0.77993 |
| *CYB5A* | AMR | rs4891541 | 0.1609126 | 0.1562776 | 0.875814 |
| *CYB5A* | AMR | rs4892193 | 0.1620219 | 0.166854 | 0.867485 |
| *CYB5A* | AMR | rs4891537 | 0.135595 | 0.065496 | 0.947779 |
| *CYB5A* | AMR | rs77442034 | 0.149931 | 1.00193 | 0.316376 |
| *CYB5A* | AMR | rs3813105 | 0.1518426 | 1.04316 | 0.296872 |
| *CYB5A* | EUR | rs72963833 | 0.04913884 | 0.30915717 | 0.75720198 |
| *CYB5A* | EUR | rs11151940 | 0.04816836 | 0.04197784 | 0.96651637 |

| | | | | | |
|---|---|---|---|---|---|
| *CYB5A* | EUR | rs72963813 | 0.0492418 | 0.25293208 | 0.8003207 |
| *CYB5A* | EUR | rs4891541 | 0.0492418 | 0.28301859 | 0.77716258 |
| *CYB5A* | EUR | rs4892193 | 0.04638716 | 0.06397648 | 0.94898896 |
| *CYB5A* | EUR | rs4891537 | 0.03633321 | 0.27166613 | 0.78587875 |
| *CYB5A* | EUR | rs77442034 | 0.04816836 | 0.49972826 | 0.61726643 |
| *CYB5A* | EUR | rs3813105 | 0.06151024 | 0.70908955 | 0.47826891 |
| *SULT2A1* | SAS | rs296360 | 0.110329 | 0.3860137 | 0.6994865 |
| *SULT2A1* | SAS | rs296361 | 0.110044 | 0.3613644 | 0.717827 |
| *SULT2A1* | EUR | rs296360 | 0.17464227 | 1.03104906 | 0.30251782 |
| *SULT2A1* | EUR | rs296361 | 0.17557699 | 0.98214294 | 0.32602944 |

**Table S6.** Population Branch Statistic (PBS) and integrated haplotype score (iHS) values for introgressed Neanderthal-derived variants across top-frequency modern human populations. PBS and iHS were computed using nf-selection. PBS p-values were assessed against the empirical genome-wide distribution; none of the variants examined fell within the top 1% of PBS values. N/A entries in iHS columns reflect the nf-selection pipeline's filtering of variants with a derived allele frequency below 5% in the target population.
